# Protection against diet-induced obesity by a single-point mutation in Kir2.1 channels

**DOI:** 10.1101/2025.09.11.675657

**Authors:** Katie M. Beverley, Marcos D. Munoz, Sang Joon Ahn, Elizabeth L. Le Master, Shane A. Phillips, Ibra S. Fancher, Pingwen Xu, Irena Levitan

## Abstract

High-fat diet (HFD)-induced obesity remains a significant global health challenge. In this study, we show that a global knock-in CRISPR mouse with the Kir2.1^L222I^ single-point mutation exhibits remarkable resistance to HFD-induced obesity. We identify palmitic acid (PA), a prevalent long-chain fatty acid in obesity, as a novel negative regulator of Kir2.1. Kir2.1^L222I^ previously shown to protect against cholesterol-mediated inhibition of Kir2.1, also confers protection against PA-induced suppression. Moreover, PA-induced suppression of Kir2.1 results in a significant loss of flow-induced vasodilation (FIV), while the L222I mutation exerts a protective effect. Notably, Kir2.1^L222I^ mice display significant protection against HFD-induced weight gain and adiposity independent of caloric intake. Specifically, the mutant mice show increased lean mass and decreased fat mass, specifically in both visceral and subcutaneous white adipose tissue (WAT) and intrascapular brown adipose tissue (BAT). Importantly, visceral-to-subcutaneous white adipose ratios decrease while BAT/WAT tissue ratios increase, suggesting a metabolically favorable fat distribution. This protection correlates with enhanced physical activity and increased energy expenditure. Metabolomic analysis reveals elevated TCA cycle metabolites in adipose tissue of Kir2.1^L222I^ mice, consistent with their enhanced energy expenditure. These findings highlight Kir2.1 channels as potential therapeutic targets for obesity and related metabolic disorders.

## Introduction

Obesity, an excess of weight and of fat deposits, is a serious, chronic and progressive disease that reached the level of a global health epidemic that affects approximately 40% adults living in the United States (1). It is a major risk factor for multiple serious conditions, including cardiovascular diseases. Obesity develops as a result of altered energy balance on a chronic basis whereby the amount of calories consumed in the diet (quantity of food or caloric content) exceeds the amount released via energy expenditure (caloric burning). This altered energy balance results in more calories being converted to and stored as fat which leads to an increase in body weight and increased adiposity (2). It is also well-established that a major contributing factor to the development of obesity is diet that is high in saturated fats and refined sugars and low in complex carbohydrates and fiber, particularly high fat Western diet (HFD) (3).

Numerous studies investigated the mechanisms of HFD-induced obesity focusing on the impacts of genetic, epigenetic and environmental factors on energy balance (4). Mechanisms regulating energy intake have been most intensely studied leading to discovery and characterization of an array of hormonal and neurologic mechanisms that regulate appetite and satiety (eg. (5–7)). In terms of energy expenditure, it is well established that its two major components are resting metabolic rate (70%) and physical activity thermogenesis (20%), the latter including spontaneous non-exercise activity and physical exercise (8). In general, obesity/decreased energy expenditure has been linked to impaired mitochondrial oxidative metabolism (eg. (9, 10)), while increased energy expenditure is associated with an increase in metabolically active tissues, such as brown adipose tissue(11) and muscle (12). Our current study discovered a novel unexpected relationship between a specific type of inwardly-rectifying K^+^ channels, Kir2.1 and propensity to HFD-induced weight gain.

Kir2.1 is an inwardly rectifying K^+^ channel ubiquitously expressed in variety of cell types. Their main physiological role is maintaining negative membrane potential of plasma membrane, which is essential for the control of ion transport across the membrane and membrane excitability (13, 14). Our group’s previous work discovered that Kir2.1 channels are essential for flow-induced activation of endothelial NO synthase (eNOS) and for flow-induced vasodilation (FIV) (15), the hallmarks of endothelial function. Our group has also previously shown that HFD-induced obesity suppresses endothelial Kir2.1 currents resulting in partial loss of FIV (16–18). We also discovered earlier that Kir2 channels, including Kir2.1, are strongly suppressed by the elevation of membrane cholesterol *in vitro* and plasma hypercholesterolemia *in vivo* (19–21) via a direct interaction between the cholesterol molecule and the channel protein (22–25). Moreover, using extensive structure-function analysis, we identified a single-point mutation, substitution of leucine in position 222 with isoleucine (Kir2.1^L222I^) that renders the channels to be cholesterol insensitive (22). Based on these studies, we generated CRISPR global knockin mouse model where wildtype Kir2.1 is globally replaced by the Kir2.1^L222I^ mutant channel (26). The model was validated to show that it renders endothelial Kir2.1 protected under hypercholesteremic conditions (26).

In this study we elucidate the mechanism of HFD-induced suppression of endothelial Kir2.1 and demonstrate that the L222I mutation confers significant protective effects against adverse changes in body composition and excessive weight gain during HFD-challenge. The findings from this investigation may provide valuable insights into potential therapeutic targets for obesity and related metabolic disorders.

## RESULTs

### Suppression of endothelial Kir currents and FIV by PA: protection by Kir2.1^L222I^ mutation

We demonstrate here that exposing endothelial cells freshly isolated from mouse mesenteric arteries to PA (50 µm, 60 min), a saturated pro-inflammatory LCFA highly prevalent in obesity (27) results in a significant (more than 2-fold) decrease in Kir currents. In contrast, exposure to Oleic Acid (OA), known for its anti-inflammatory properties (28) has no effect under the same experimental conditions. This is shown by typical whole cell Kir current traces recorded in response to voltage ramps from single cells (the shape of the current curves reflects the typical current-voltage relationship of Kir2 channels) (Fig.1A) and by Kir current densities at 100 mV (Fig.1C). Furthermore, using a Kir2.1^L222I^ CRISPR mouse model (global substitution of Kir2.1 for Kir2.1^L222I^ developed on the *C57BL6J* genetic background) we found that *Kir2.1^L222I^* mutation confers full protection to the endothelial Kir2.1 currents against PA-induced suppression under these experimental conditions (Figs.1B,C).

**Figure 1.**
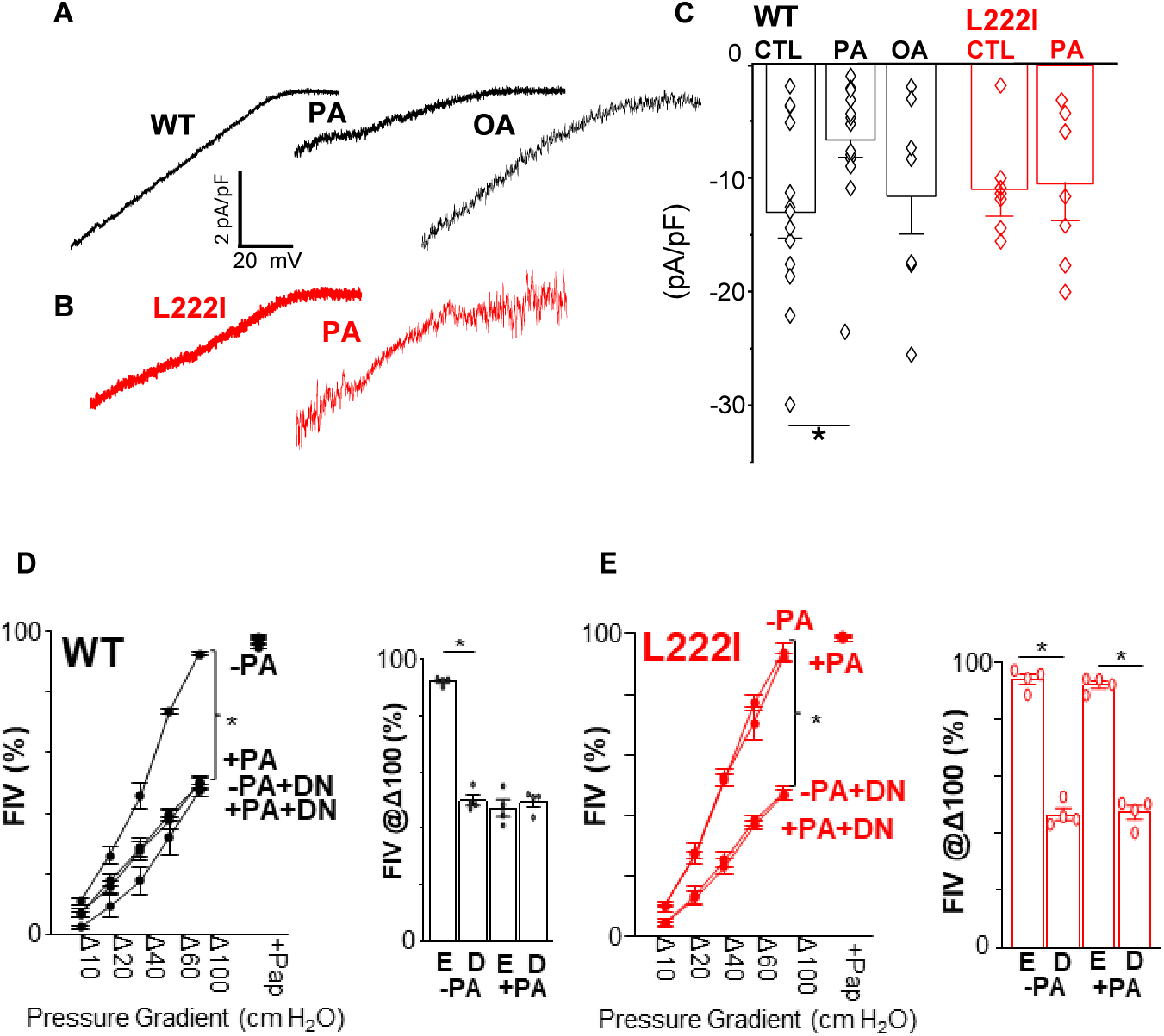
PA suppresses endothelial Kir2.1^WT^ but not Kir2.1^L222I^. A,B: Representative current traces recorded from mesenteric endothelial cells freshly isolated from Kir2.1^WT^ or Kir2.1^L222I^ mice exposed to 50 mM PA or OA for 1 hour vs. control. C: Current Densities (pA/pF) from control and PA treated (n=8,14 (p=0.0135) Kir2.1^WT^ and Kir2.1^L222I^ (n=8,9 p=0.518) from 5 independent experiments. D and E: FIV in mesenteric arteries from Kir2.1^WT^ or Kir2.1^L222I^ in the absence and presence of PA and comparison at Δ100 cm H_2_O. (n=4 Kir2.1^WT^, n=4 Kir2.1^L222I^, *p<0.05) E=Empty Adv viral vector, D=Kir2.1dominant negative expressing Adv viral vector.

Building on these findings, we next examined the functional implications of PA-induced suppression of Kir2.1. We also show here that exposing freshly-isolated mesenteric arteries to PA results in significant suppression of FIV (see the flow dose response curves and maximal dilation at 100 cm H_2_O), with no further decrease when the arteries are transduced with a dominant-negative Kir2.1 construct driven by an endothelial specific promoter Cdh5 (Fig. 1D) indicating that PA-induced impairment of FIV can be attributed to the loss of endothelial Kir2.1 function. Furthermore, we also show here that Kir2.1^L222I^ mice have full vasodilation in the presence of the PA challenge, similar to controls (Fig. 1D). Our data also show that FIV in Kir2.1^L222I^ arteries exposed to PA remains sensitive to dnKir2.1 indicating that the activity of the channels in these mice is intact. There was no difference in vasodilation in response to flow between Kir2.1^WT^ and Kir2.1^L222I^ mice in control conditions (Fig. 1E). Under control conditions, the FIV in both strains was also equally sensitive to the downregulation of the endothelial Kir2.1 by dnKir2.1.

### Protection of endothelial Kir currents and FIV against obesity-induced damage by Kir2.1^L222I^ mutation

To determine whether the Kir2.1^L222I^ mutation also confers protection to endothelial Kir2.1 currents and FIV *in vivo* in a mouse model of HFD-induced obesity, Kir2.1^WT^ and Kir2.1^L222I^ mice were either fed a High Fat Diet (42% fat, 34% sucrose) for 8-10 weeks starting at 8 weeks of age or maintained on a regular low-fat diet (LFD) for the same period. Currents were recorded from freshly isolated mesenteric endothelial cells on the day of the isolation. As we have shown previously (16), feeding Kir2.1^WT^ mice HFD virtually eliminates Kir2.1 activity in mesenteric endothelial cells, as evident in the representative current traces (Fig.2A) and current densities at -100 mV (Fig. 2C). Our new data show that Kir currents in Kir2.1^L222I^ and Kir2.1^WT^ mice were similar under LFD but not under HFD conditions. In contrast to Kir2.1^WT^ currents, diet-induced obesity had no significant effect on Kir2.1^L222I^ currents (Figs 2B,C). These data show that Kir2.1^L222I^ currents are significantly larger than Kir2.1^WT^ currents under HFD challenge, indicating that this mutation confers significant protection to Kir2.1 currents.

**Figure 2.**
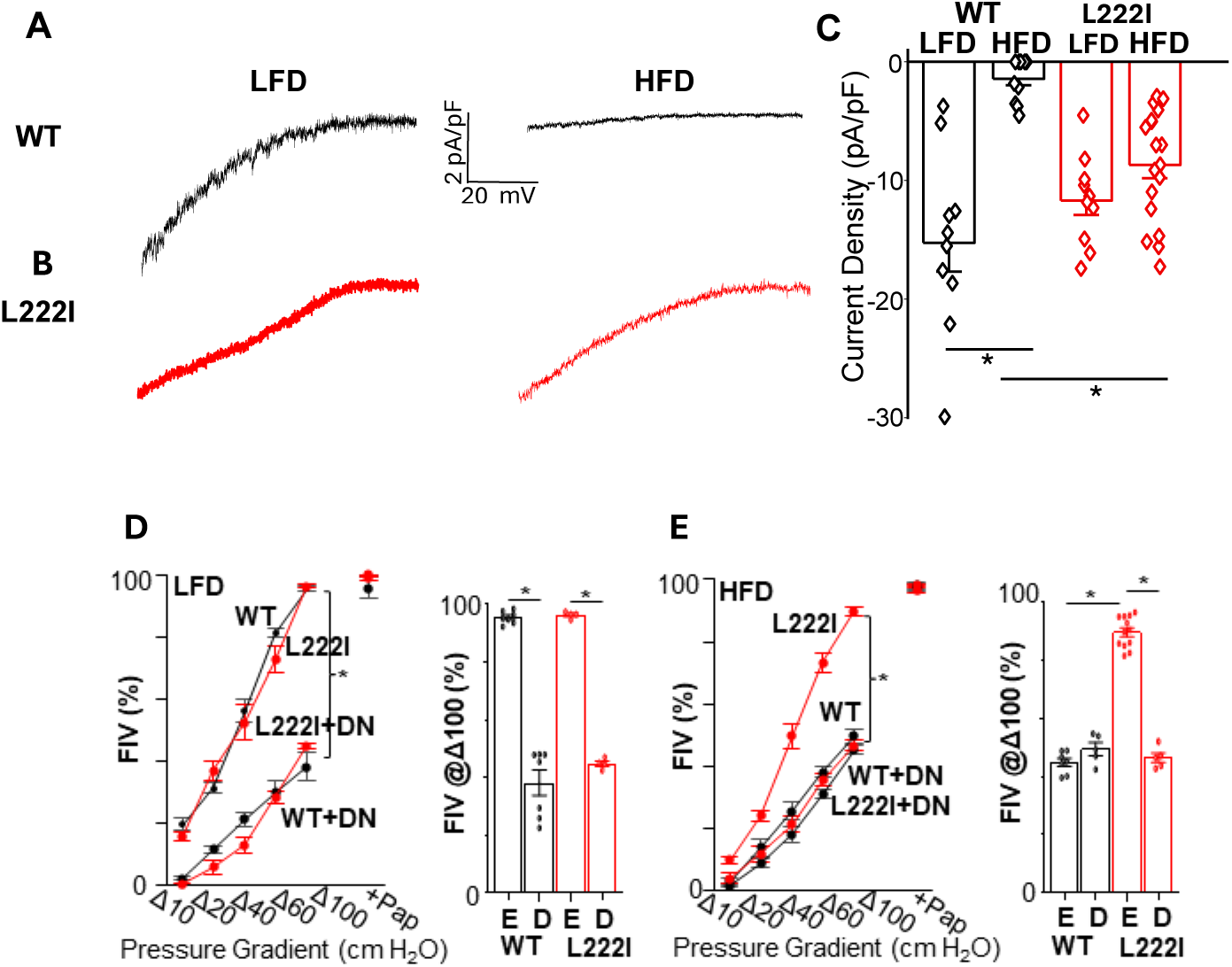
Kir2.1^L222I^ is functionally protective against HFD induced suppression. Representative current traces from Kir2.1^WT^ (A) and Kir2.1^L222I^ (B) freshly isolated endothelial cells from mice on LFD or HFD. C: Current Densities (pA/pF) at -100 mV from individual recordings. Kir2.1 WT current density was suppressed by the change from LFD to HFD (p<0.0001, n=10,11 cells recorded from 8 mice per group). Kir2.1 ^L222I^ current density was unaffected by the change in diet (p=0.25, n=10,18 cells recorded from 8 and 12 mice per group respectively). Under HFD conditions Kir2.1^WT^ had significantly reduced current density compared with Kir2.1^L222I^ (p=0.0006). C:FIV in mesenteric arteries from low-fat diet fed mice and comparison at Δ100 cm H_2_O. (n=7 WT, n=4 Kir2.1^L222I^, *p<0.05) D: FIV in mesenteric arteries from high-fat diet fed mice and comparison at Δ100 cm H_2_O. (n=7 WT, n=12 Kir2.1^L222I^, n=5 for WT and Kir2.1^L222I^ with DN, *p<0.05) E=Empty Adv viral vector, D=Kir2.1dominant negative expressing Adv viral vector.

As the Kir2.1-dependent component of FIV has previously been shown to be suppressed by HFD (16), we next evaluated whether the protective effects of the Kir2.1^L222I^ mutation on Kir2.1 currents translated to preserved vasodilation responses. Kir2.1^WT^ and Kir2.1^L222I^ mice were fed either LFD or HFD using the same diet regimen as above, after which mesenteric arteries were isolated for FIV analysis. Under LFD conditions, we observed no difference in FIV between Kir2.1^WT^ and Kir2.1^L222I^ mice (Fig. 2D).

However, under HFD conditions, a significant difference emerged: FIV in Kir2.1^WT^ arteries in mice fed HFD was significantly lower than in Kir2.1^L222I^ arteries (Fig. 2E). Similarly to PA, transducing the arteries with endothelial-specific dnKir2.1 decreased FIV in both Kir2.1^L222I^ and Kir2.1^WT^ under LFD but only in Kir2.1^L222I^ on HFD (Fig.2D,E). These findings indicate that the Kir2.1^L222I^ mutation is protective against HFD-induced suppression of the Kir2.1 component of endothelial FIV. Noteworthy, the HFD-induced effects on endothelial Kir2.1 and FIV mirror those of the PA exposure.

### No differences in feeding behavior between Kir2.1^WT^ and Kir2.1^L222I^ mice

Next, we tested whether the protective effect of Kir2.1^L222I^ mutation could be due to differences in the feeding behavior: if Kir2.1^L222I^ mice consume less food, when on HFD, that could potentially lead to the observed protective effect on the channel function and FIV. We found, however, that this is not the case. The feeding behavior was evaluated using a BIODAQ system that continuously monitors the weight of food consumed by individual mice, as well as the number, duration, and frequency of interactions with the food hopper (29).

Our study included two independent cohorts of age-matched Kir2.1^WT^ and Kir2.1^L222I^ mice after 14 weeks on HFD (4-5 animals per group in each cohort). The animals were acclimated for 3 days in the BIODAQ system and then monitored continuously for 6 days while receiving the same HFD diet as prior to the experiment.

All experimental groups of animals consumed the same number of meals per 24h for the duration of the study showing successful acclimation to the system (Fig.3A). No significant differences were observed between the two strains in the size of individual meals (Fig. 3B) or the number of meals per day (Fig. 3C). When examining day/night meal distribution, all mice exhibited typical nocturnal behavior with more food consumed during the night than day. This pattern is illustrated by representative histograms of food consumption for individual Kir2.1^WT^ and Kir2.1^L222I^ mice (Fig. 3D,E) and the average number of meals consumed during 12h periods of day (7AM-7PM) and night (7PM-7AM) following the light cycle in the experimental system (Fig. 3F). While both Kir2.1^WT^ and Kir2.1^L222I^ mice took less meals during the day compared to night, we did observe that Kir2.1^L222I^ mice had more daytime meals than Kir2.1^WT^ mice. However, this did not result in a significant increase in the average number of meals per 24h because there was no difference between the two strains during the nighttime feeding, when the majority of consumption occurred.

**Figure 3.**
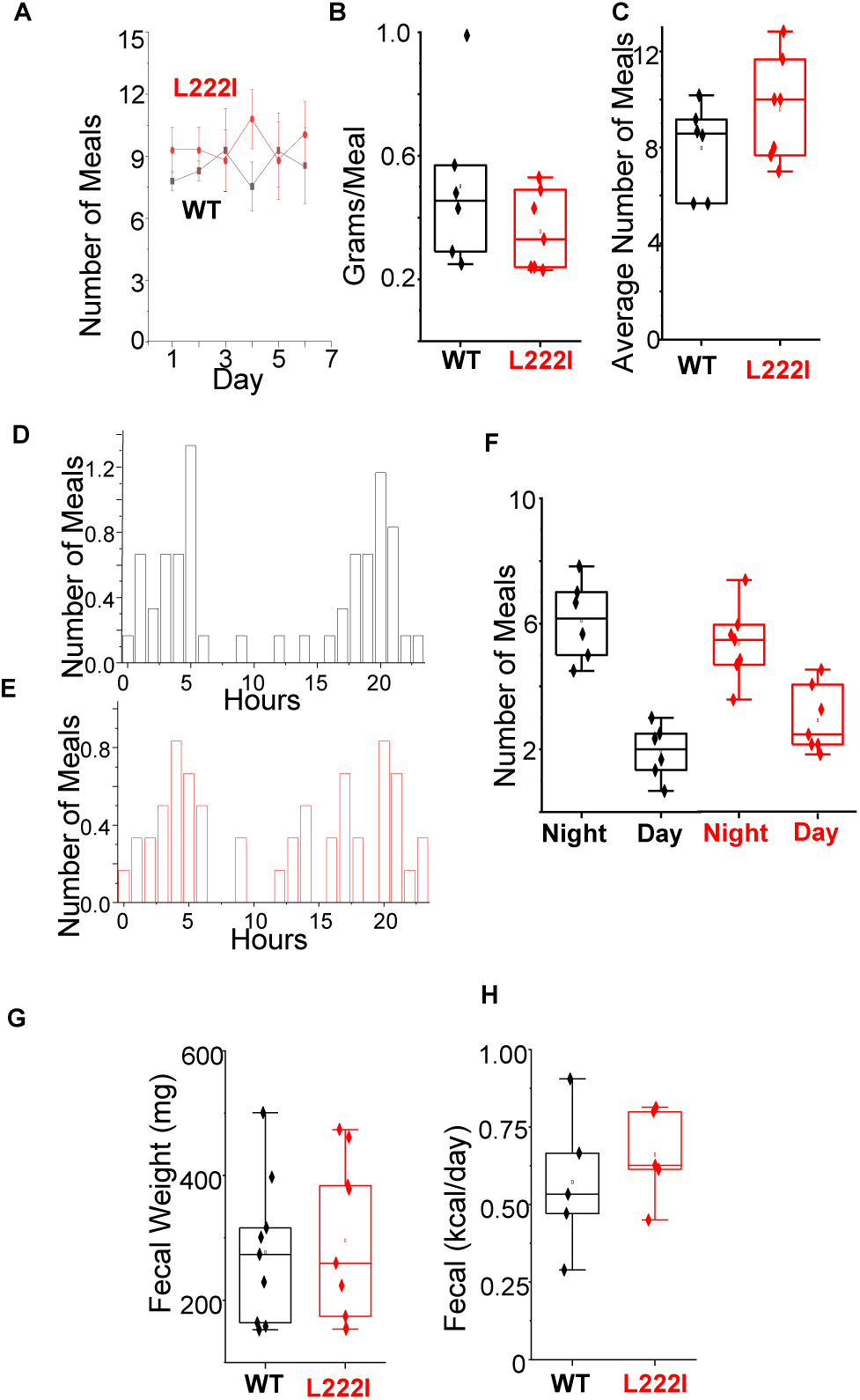
Kir2.1^L222I^ mice and Kir2.1^WT^ have the same feeding behavior and fecal content. A: Average meals over study time course for Kir2.1^WT^ (n=5) and Kir2.1^L222I^ (n=7) mice. B:The average size of the meals and average number of daily meals (C). Average meal distribution over 24 hours for a single Kir2.1^WT^ (D) or Kir2.1^L222I^ (E) mouse. F: Number of day and night meals per mouse. G-H: There was no difference in total fecal mass (n=8/9) or fecal caloric content (n=5).

### No differences in fecal caloric content

Following our feeding evaluation, we investigated whether the Kir2.1^L222I^ mutation affects fecal excretion and caloric content. We used a Techniplast metabolic cage system featuring a specialized funnel design that separately collects feces and urine, preventing cross-contamination (30). For our analysis, we used age-matched Kir2.1^WT^ and Kir2.1^L222I^ mice maintained on HFD for 14 weeks. The study comprised two independent cohorts for fecal weight measurements and one cohort for BOMB calorimetry (4-5 animals per group in each cohort). BOMB calorimetry measures the heat produced by burning fecal pellets to determine total kilocalories. We conducted experiments in pairs (2 Kir2.1^WT^ and 2 Kir2.1^L222I^) running simultaneously. Each animal was placed individually in a Techniplast cages for 48 hours. Our results showed no significant difference between the two groups in either total fecal mass per mouse over the 48-hour period (Fig. 3G) or kilocalories excreted in feces per day (Fig. 3H).

### Protective effects of Kir2.1^L222I^ against high-fat diet-induced body weight gain

Unexpectedly, the Kir2.1L222I mutation demonstrates a protective effect against diet-induced obesity, despite having no impact on feeding efficiency. When subjected to a 14-week high-fat, high-caloric diet regimen, Kir2.1^L222I^ mice showed remarkable resistance to weight gain compared to Kir2.1^WT^ mice. The differences in body weight emerged within the first week of HFD and progressively increased throughout the study (Fig. 4A). By the study’s conclusion, Kir2.1^WT^ mice had gained approximately 15% more body weight on HFD compared to LFD, while Kir2.1^L222I^ mice maintained similar weights regardless of diet type (Fig.4B). Importantly, this divergence was specific to the HFD, as no differences were observed between the groups fed LFD (Fig. 4B). As reported previously (31), female C57BL/6 mice gained significantly less weight than males across all conditions, with no genotype-specific differences observed (*Suppl. Fig. 1*).

**Figure 4.**
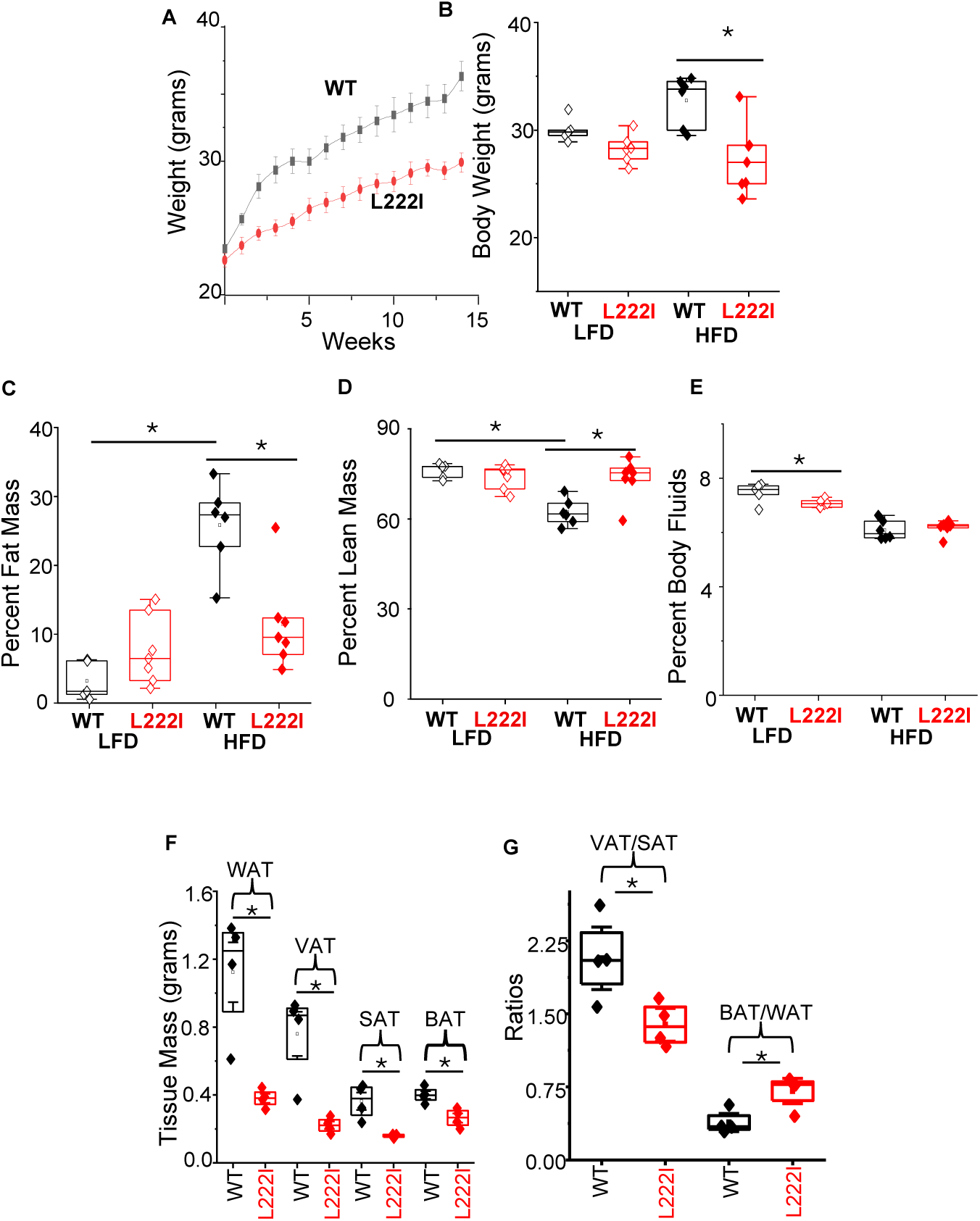
Kir2.1^L222I^ mice are protected against HFD induced changes in body composition. A: Time course of body weights of Kir2.1^WT^ (n=9) and Kir2.1^L222I^ (n=10) mice on HFD for 14 weeks. B: Body weights of mice after 14 weeks fed either LFD or HFD (*p<0.001, n=5-7 mice per group). C-E: NMR of %body fat mass, %lean mass, and %body fluids respectively of Kir2.1^WT^ and Kir2.1^L222I^ fed LFD or HFD measured at the end of the 14 week on HFD (*p<0.001, n=5-7 mice per group). F-G: Mass of fat visceral, subcutaneous, and brown adipose depots from Kir2.1^WT^ and Kir2.1^L222I^ mice (n=4 mice per group, *p<0.001).

Beyond weight differences, the Kir2.1^L222I^ mutation also conferred beneficial effects on body composition, as assessed by NMR analysis. After 14 weeks of HFD, Kir2.1^WT^ mice showed an approximately fourfold increase in fat mass, whereas Kir2.1^L222I^ mice maintained stable fat mass while showing a ∼15% increase in lean mass (Figs.4C,D). No difference was observed in %body fluids between Kir2.1^WT^ and Kir2.1^L222I^ (Fig.4E). Anatomical analysis further confirmed these findings, with Kir2.1^WT^ mice developing significantly larger fat depots compared to Kir2.1^L222I^ mice (Fig. 4F). Additionally, Kir2.1^L222I^ mice displayed metabolically favorable fat distribution, characterized by higher brown adipose tissue to white adipose tissue (BAT/WAT) ratio and lower visceral adipose tissue to subcutaneous adipose tissue (VAT/SAT) ratio (Fig. 4G).

Serum lipid analyses further supported the protective effects of the Kir2.1^L222I^ mutation. While both genotypes on HFD exhibited elevated total cholesterol and low-density lipoprotein (LDL) levels compared to those on LFD, these increases were significantly attenuated in Kir2.1^L222I^ mice (Suppl. Fig. 2). Similarly, high-density lipoprotein (HDL) levels increased in both strains on HFD, but to a lesser extent in Kir2.1^L222I^ mice. No differences were observed in LDL/HDL ratios or triglyceride levels between the two strains on either diet. Collectively, these results support a protective role for the Kir2.1^L222I^ mutation in the adverse effects of HFD on body weight, composition, and lipid metabolism.

### Kir2.1^L222I^ mutation results in increased RER and running wheel physical activity

Notably, we found that protection against HFD-induced obesity in Kir2.1^L222I^ mice is associated with an increase in Respiratory Exchange Rate (RER = CO2/VO2), an indirect indicator of metabolic fuel consumption. Using the Iworx metabolic cage system, we monitored oxygen and carbon dioxide in a closed system from individual mouse on an hourly basis over a 24 hour period. This experiment was conducted on Kir2.1^WT^ and Kir2.1^L222I^ mice fed HFD for 14 weeks across two independent cohorts with 8-10 mice per group.

Our data demonstrates that Kir2.1^L222I^ mice have a significantly higher RER compared with Kir2.1^WT^ (Fig. 5A). Specifically, the average RER in Kir2.1^WT^ mice on HFD was ∼0.75, indicating primarily lipid oxidation as a fuel source, while Kir2.1^L222I^ mice had an RER of ∼0.85, suggesting a higher proportion of carbohydrate metabolism. In consistency with this metabolic shift toward carbohydrate utilization, we observed a significant decrease in glucose level in serum of Kir2.1^L222I^ mice compared with Kir2.1^WT^ (Fig. 5B).

**Figure 5.**
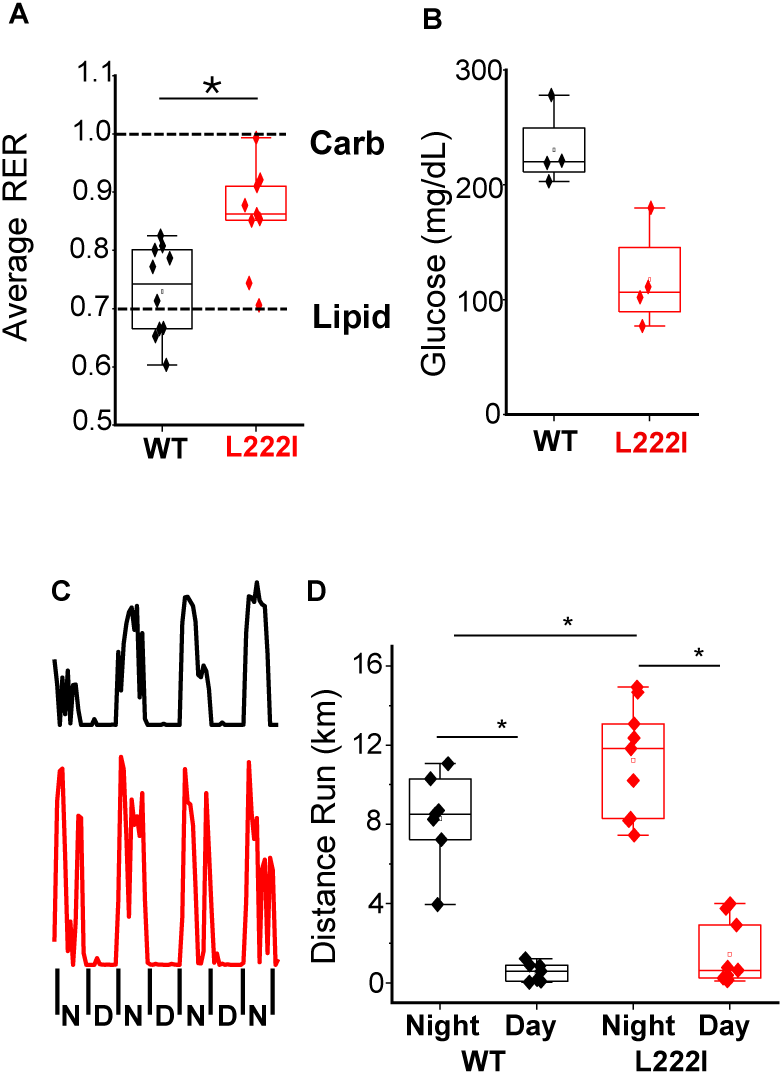
Kir2.1^L222I^ mice have increased RER, decreased serum glucose, and physical activity on the running wheel. A: Average RER for Kir2.1^WT^ and Kir2.1^L222I^ mice on HFD (n=9, p=0.02). Carbohydrates (carbs) and lipids are denoted at 1.0 and 0.7 respectively, indicating their use as a primary fuel source. B: Serum glucose levels from Kir2.1^WT^ and Kir2.1^L222I^ mice after 16 weeks on HFD. C: Representative running wheel movement traces from Kir2.1^WT^ and Kir2.1^L222I^ mice on HFD. D: Average distances run during the day and at night for both Kir2.1^WT^ (n=7) and Kir2.1^L222I^ (n=9) animals (*p<0.01).

Furthermore, Kir2.1^L222I^ mice have enhanced physical activity in running wheel experiments. These mice ran significantly more meters on average than Kir2.1^WT^ mice during the night active period (Fig. 5C), with representative activity traces shown in Fig. 5D. Taken together, these findings indicate that both altered metabolic fuel preference and increased physical activity potentially contribute to the differences in body composition between the two mouse strains.

### Metabolic profile of Kir2.1^L222I^ mice at early stages of HFD exposure

Our findings suggest that reduced body weight gain in Kir2.1^L222I^ mice may stem from metabolic adaptations to HFD. To explore this hypothesis, we conducted indirect calorimetry studies using the Promethion System (Sable), which provides real-time measurements of energy expenditure (EE), RER, physical activity, and food/water intake. Eight-week-old male Kir2.1^WT^ and Kir2.1^L222I^ mice were acclimated to the system for one day, followed by a three-day baseline period on a LF chow diet and then three days on an HFD. Energy expenditure was compared between the groups using ANCOVA analysis to account for body weight differences (32).

Interestingly, we found that Kir2.1^L222I^ mice exhibited significantly higher EE than WT mice during the chow diet period (Fig. 6A), with a strong trend toward increased EE on the HFD (P=0.06, Fig. 6B). This heightened EE corresponded closely with increased physical activity in Kir2.1^L222I^ mice, as measured by home cage voluntary locomotion, which remained consistently higher than in WT mice across both chow and HFD conditions (Fig. 6C). These results suggest that enhanced energy metabolism and activity levels may contribute to the protective effects observed in Kir2.1^L222I^ mice.

**Figure 6.**
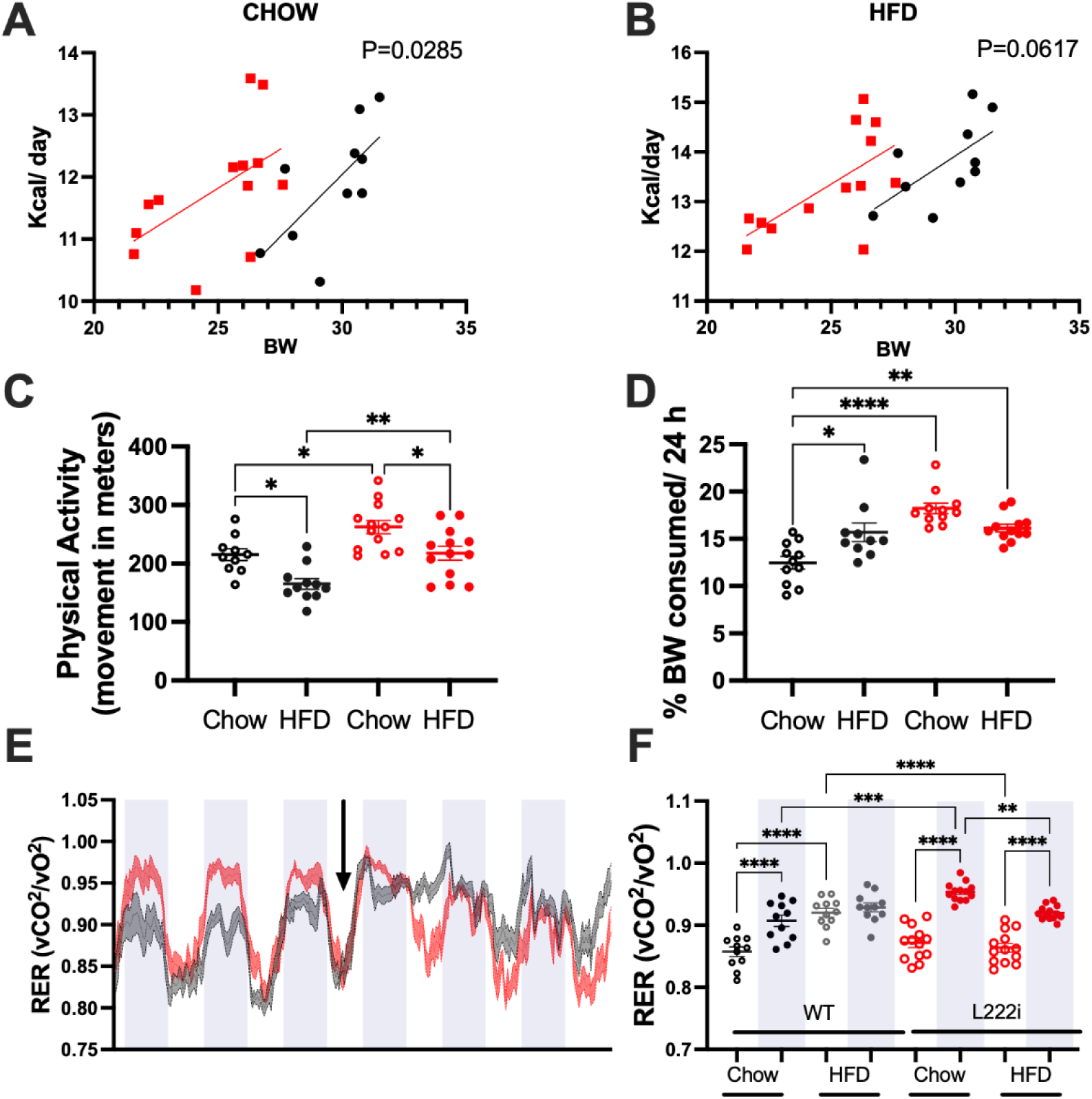
Metabolic Phenotyping at Early Time-Point. A-B: ANCOVA analysis of energy expenditure with bodyweight as covariate for Kir2.1WT and Kir2.1L222I mice on chow diet (A) and HFD diet (B) periods. C: Physical activity expressed as translational movement of mice in meters. D: Average food intake normalized by mouse bodyweight. E: Continuous measurement of RER (v02/vCO2) through the 7 days of experiment. Black arrow indicates the time of diet change from chow to HFD diet. F: Averages of RER values for Kir2.1WT and Kir2.1L222I mice on day/night period. Values are expressed as averages of 72 hours on each diet period in A-D and F. On E-F White/grey dashes indicate day and night period respectively.

Further supporting this metabolic advantage, we observed distinct differences in substrate utilization. When switched to an HFD, WT mice displayed reduced circadian oscillation in the RER, indicating disrupted substrate utilization (Fig. 6E). In contrast, Kir2.1^L222I^ mice maintained robust RER oscillations during the HFD period (Fig. 6F), suggestive of a healthier metabolic profile. Additionally, the elevated EE and physical activity in Kir2.1^L222I^ mice aligned with a higher RER during the night on a chow diet compared to WT mice. These findings on EE, RER, and physical activity are consistent with our earlier observations that WT mice accumulate fat mass on HFD, while Kir2.1^L222I^ mice are protected from this effect (Fig. 4C).

To complete the picture of metabolic regulation, we examined food intake. Our study on mice fed an HFD for 14 weeks showed no differences in feeding behavior between Kir2.1^L222I^ and WT mice (Fig. 3A-E). However, Promethion System data revealed that Kir2.1^L222I^ mice consumed approximately 5% more food (BW-normalized) than WT mice on a chow diet. Remarkably, Kir2.1^L222I^ mice did not increase food intake when transitioned to an HFD, while sustaining higher EE (Fig. 6D). These results indicate a well-balanced energy equation (food intake versus EE) in Kir2.1^L222I^ mice on a chow diet, which aligns with their lower body weight on HFD, driven by stable food intake and elevated EE.

Collectively, these findings underscore a role for the Kir2.1^L222I^ mutation in promoting metabolic resilience against HFD-induced weight gain and fat accumulation.

### An increase in TCA cycle metabolites in Kir2.1^L222I^ mice

We next performed metabolomics analysis on mesenteric white adipose tissue from Kir2.1^WT^ and Kir2.1^L222I^ mice (4-6 animals per group) after 14 weeks on HFD. Our aim was to investigate metabolic differences that might explain the increased energy expenditure. The metabolomics analysis focused on the primary components of cellular metabolism: fatty acids, carbohydrates, carnitines, and amino acids (33). From both mouse groups, we identified 203 unique metabolites. As illustrated in the volcano plot (Fig. 7A), the majority of these metabolites showed increased levels in Kir2.1^L222I^ mice compared to Kir2.1^WT^ controls. In particular, we observed significant upregulation of the TCA/citric acid cycle, a central metabolic pathway for cellular energy production. This upregulation is demonstrated in the heatmap of the TCA cycle metabolites (Fig. 7B). Specifically, we detected elevated levels of the high-energy electron carrier flavin adenine dinucleotide (FAD) along with increases in three cycle intermediates: fumarate, malate, and citrate (Fig. 7B).

**Figure 7.**
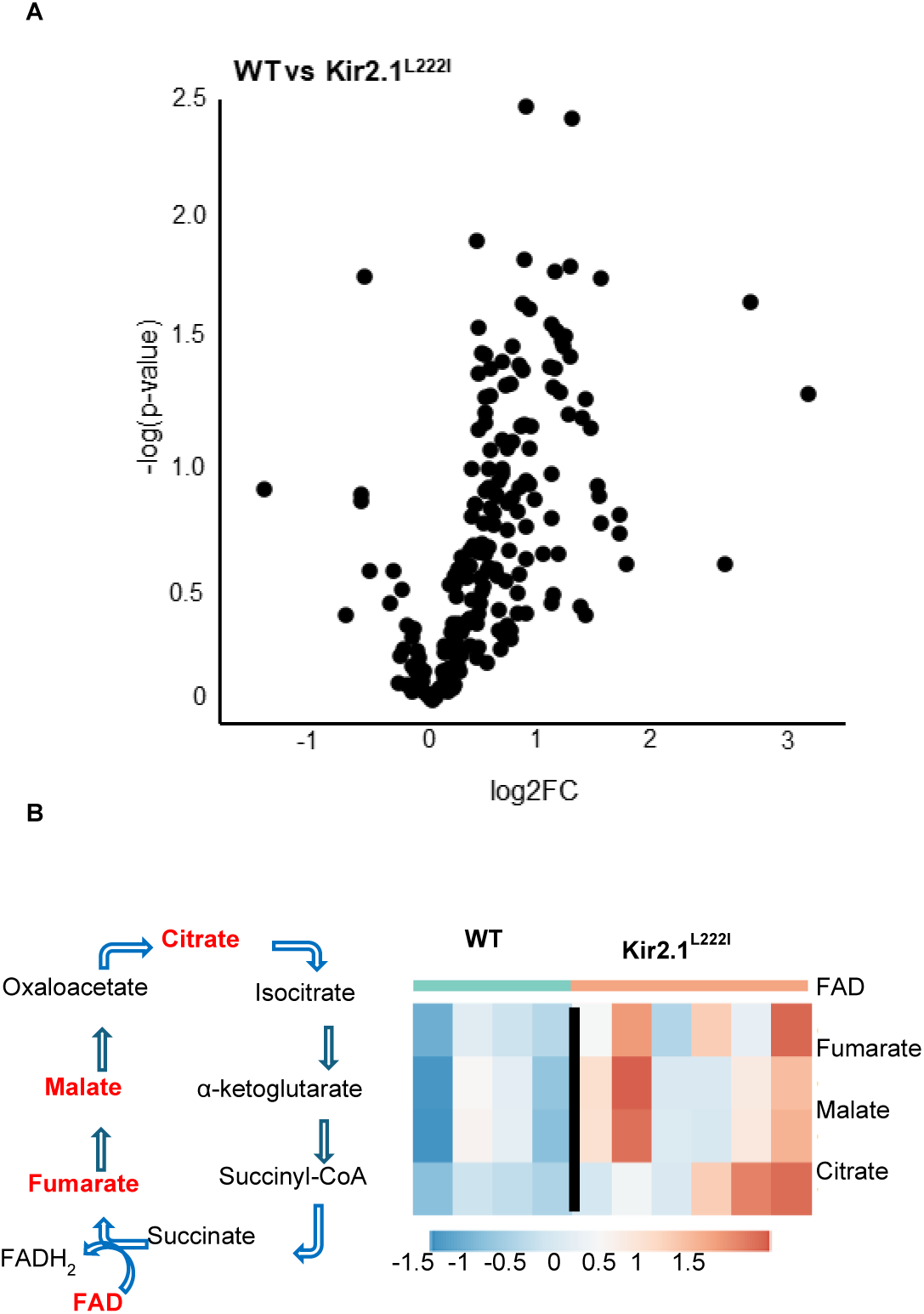
Metabolomic analysis of mesentery from Kir2.1^WT^ and Kir2.1^L222I^ suggesting changes in TCA Cycle. A: Volcano Plot of metabolites present in mesenteric adipose tissues from Kir2.1^WT^ and Kir2.1^L222I^ mice from metabolomic analysis. B:Schematics of TCA/Citric Acid cycle with identified metabolites highlighted in red and heatmap analysis from mesenteric adipose tissues.

## Discussion

In this study, we discover that gene editing of a major type of K+ channels, Kir2.1, ubiquitously expressed in a variety of tissues, confers significant protection against HFD-induced obesity in mice. We also show that this protection effect correlates with improved endothelial function and increased movement of the animals, suggesting that this might contribute to improved energy balance. First, we show that Kir2.1 channels are suppressed by a major lipid component of HFD, PA (34), indicating that PA may be responsible for the HFD-induced loss of Kir2.1 function. Moreover, we found that a mutation that we have shown earlier to render the channels to be insensitive to cholesterol (22), also renders them to be insensitive to PA, suggesting that cholesterol and PA suppress the channels by a similar mechanism. As expected (15), the rescue of endothelial Kir2.1 function leads to the rescue of FIV. Notably, congenital global Kir2.1 ^L222I^ mice are protected from HFD-induced weight gain and associated changes in body composition, associated with increased energy expenditure and physical activity. This is the first study to demonstrate that the energy expenditure in HFD-induced obesity depends on Kir channels.

Our earlier studies showed that Kir2.1 channels in the endothelium of resistance arteries are suppressed by HFD-induced obesity, leading to a significant impairment of FIV (16). While we initially hypothesized these effects are mediated by cholesterol (a known Kir2.1 suppressor), we subsequently discovered that HFD-induced suppression of FIV is cholesterol independent (16). Here, we identify PA, a prevalent pro-inflammatory long-chain fatty acid (LCFA) accumulated in obesity, as a critical factor suppressing both Kir2.1 channels and FIV. Mechanistically, little is known about the regulation of Kir channels by LCFAs, particularly by PA. Several earlier studies focused on *oleoyl-coA,* an ester of oleic acid with coenzyme A, a metabolic intermediate elevated in obesity (35) and found that it suppresses Kir2.1 channels, as well as Kir1.1, Kir3.4, and Kir7.1 (36) but to enhance K-ATP (Kir6.2) (37). These effects were attributed to either competition with or enhancement of the regulatory phospholipid PI(4.5)P_2_ (PIP2) (36, 37), which is essential for Kir function (38). Our study discovers that the same single-point mutation abrogates the sensitivity of Kir2.1 to cholesterol and to PA suggesting that the two inhibitory effects share a similar mechanism. The mechanisms of cholesterol-induced suppression of Kir2.1 have been thoroughly studied by our group (19, 22, 24, 25, 39, 40). Briefly, we showed that cholesterol suppresses Kir2 channels by direct binding to hydrophobic pockets in the transmembrane domains (25), which are adjacent to known PIP2 binding sites (40), consistent with the disruption of Kir2.1-PIP2 interaction. The residue leucine L222, which we find critical for the PA-suppression of Kir2.1, was shown earlier to be important for both cholesterol and PIP2 sensitivities of the channels (22, 41). Thus, we postulate that PA also suppressed Kir2.1 channels by interfering with PIP2-dependent gating.

In terms of the connection between lipid-induced suppression of Kir2.1 and FIV, our previous studies demonstrated that over-expression of Kir2.1 in the endothelium of resistance arteries leads to full recovery of FIV in both hypercholesterolemia and obesity, in mice and humans. Here, we show that the Kir2.1^L222I^ mutation fully prevents HFD-induced decrease in FIV, suggesting improved endothelial and cardiovascular function under HFD challenge. Our further studies demonstrate that this improvement of endothelial function correlates with improved energy balance.

We also discover in this study that rendering Kir2.1 channels to be lipid (PA/cholesterol) insensitive leads to global protection against HFD-induced metabolic challenges. Previous studies found that two types of Kir channels, Kir6.2 and Kir7.1 are involved in the regulation of feeding behavior and satiety (42, 43). Specifically, Kir6.2 in the hypothalamus is suppressed by leptin (44), an adipocytes-derived hormone signaling satiety to the brain (7). Kir7.1 has also been shown to promote food intake, where activation of Kir7.1 by MC4R signaling pathways in the choroid plexus leads to increased neuronal firing and hunger signals (45). Additionally, both Kir6.2 and Kir2.1 play important roles in insulin secretion from pancreatic beta cells and subsequent glucose uptake, as extensively documented in diabetes research (46). Our studies are unique in demonstrating that lipid-induced suppression of Kir2.1 plays an important role in the regulation of energy expenditure. This conclusion is based on several lines of evidence: (i) Kir2.1^L222I^ mice do not gain weight or experience diet-induced changes in body composition under HFD challenge, despite consuming the same amount of food and having identical fecal caloric excretion as Kir2.1^WT^ animals that develop obesity, (ii) Kir2.1^L222I^ mice have an increase in energy expenditure under both LFD and HFD conditions (iii) Kir2.1 ^L222I^ exhibit an increase in voluntary wheel running and home-cage locomotive activity, contributing to higher overall energy expenditure.

Another interesting observation is that Kir2.1^L222I^ mice maintain consistently elevated RER, showing preferential carbohydrate usage over fat oxidation during both LFD feeding and chronic HFD challenges. While Kir2.1^WT^ mice acutely develop disrupted circadian RER patterns on HFD, mutant mice maintain stable substrate utilization. This metabolic stability likely stems from a hypermetabolic state characterized by increased energy expenditure and physical activity, potentially involving enhanced thermogenesis in fat tissues. This state drives rapid carbohydrate burning to fuel heat production. The resulting elevated energy expenditure depletes calories, prevents positive energy balance, and protects against the weight gain and metabolic dysfunction typically caused by HFD.

Mechanistically, we analyzed the metabolomics profiles from mesenteric adipose of Kir2.1^WT^ and Kir2.1^L222I^ mice. We observed upregulation of the TCA/citric acid cycle in Kir2.1^L222I^ tissues, a pathway typically suppressed under obese conditions in both mice and humans, particularly during insulin insensitivity (47–49). Specifically, we detected increased levels of TCA/citric acid cycle intermediates: FAD, fumarate, malate, and citrate. Elevated FAD, a high-energy electron carrier, suggest enhanced metabolic efficiency under higher metabolic demands, such as during increased physical activity (50). Similarly, higher levels of fumarate, malate, and citrate indicate increased energy production (ATP) during activity (51). The role of Kir2.1 function in the metabolic pathways is not yet understood but its critical role in vascular health suggests a potential for an enhanced role in activity consistent with these findings.

In summary, in this study we show that when the Kir2.1 channel is rendered insensitive to suppressive lipids with the Kir2.1^L222I^ mutation the animals have increased activity which may contribute to their protection against diet induced obesity. We propose that this increased activity likely results from preserved endothelial function, given the well-established link between cardiovascular function and physical activity (52). When vascular function becomes impaired, physical activity typically decreases due to reduced tissue oxygenation, significantly affecting overall activity levels (53).

A limitation of this study is that the Kir2.1^L222I^ animal model is a congenital global knock-in of the mutation across all cells, making it difficult to determine which specific tissue drives the observed protection against HFD-induced metabolic challenge. While our data show protected endothelial function in these mice, this model cannot definitively determine whether endothelial function alone is responsible for the systemic protection against diet-induced changes in body composition. In light of the altered circadian rhythm we observed in the feeding behavior it is possible that Kir2.1 in the brain is contributing to this protection against obesity. It is also possible that increased insulin secretion from pancreatic beta cells, which are known to express Kir2.1 (46), that improves overall glucose sensitivity and protects against the development of obesity associated diabetes contributes. It remains unknown whether adipocytes themselves express Kir2.1, though adipocyte-associated macrophages do express Kir2.1 (54) and could potentially drive the observed changes in adiposity. Future studies using tissue-specific knock-in of the Kir2.1^L222I^ mutation would allow us to determine which tissue primarily drives the systemic protection against diet-induced changes that we observed.

## Methods

### Animal Models and Diets

In this study, we used C57BL6J male and female mice. We also previously generated the Kir2.1^L222I^ global CRISPR knock-in mouse model (26) which was also utilized in the study. Mice were either maintained on chow or given Western-style diet (TD.88137, from Inotiv) for a consistent specification of 21% milk fat, 34.5% sucrose, and 0.2% cholesterol (by weight) beginning at 8 weeks of age. Animals were maintained on diet for 8 weeks (Patch-Clamp, FIV, Running Wheels, Promethion) or 14 weeks (weights, NMR, Biodaq, Techniplast, iWorx).

### Patch Clamp Electrophysiology

To assess the effects of long-chain fatty acids on Kir2.1 channel currents and to evaluate the effects of the Kir2.1^L222I^ mutation on currents under HFD challenge we performed patch clamp electrophysiology on freshly isolated endothelial cells from mouse mesenteric arteries. Mouse mesenteric arteries were harvested from 16-wk-oldC57BL6J mice; Kir2.1^WT^ and Kir2.1^L222I^ after 8 weeks of chow diet or HFD. ECs were then isolated from these arteries for patch-clamp recording as previously described (16, 26, 55). Freshly isolated cells in suspension were allowed to adhere directly onto the glass coverslip and were subject to control, PA , or OA (50 µM for 1 hour) treatment.

Pipettes with resistance (2–4 MΩ) were pulled using borosilicate glass (BF150-11010, Sutter) and a vertical puller (Model PP-830). Currents were recorded using an EPC10 amplifier and the PatchMaster Next (HEKA Electronik), filtered at 2 kHz, and digitized at 10 kHz. ECs were held at -30 mV and a voltage ramp of -140 to +40 mV was applied over 400ms. The perforated patch was obtained by adding amphotericin B (250 μg/mL) to the pipette solution. When necessary, leak subtraction was performed offline to collect the most accurate data points at -100 mV for analysis. Patch solutions were the following: Extracellular solution contained (in mM): 90 NaCL, 60 KCl, 10 HEPES, 1.5 CaCl2, 1 MgCl2, 1 EGTA, pH 7.3. Pipette solution contained (in mM): 145 KCl, 10 HEPES, 1 MgCl2, 4 ATP, 1 EGTA, pH 7.3 (26).

### Flow-induced vasodilation

To investigate the role of Kir2.1^L222I^ in vascular function under obesity or PA, mouse mesenteric arteries were used and measured flow-induced vasodilation in ex vivo using flow chamber and pressure gradient induced flow as previously described (15, 26). Briefly, 75 – 200 μm diameter mesenteric arteries were cannulated in between the two pulled glass pipettes. Two connected reservoirs to each glass pipette were filled up with Krebs buffer and located at 60 cm for 30 minutes to rest the vessel in 37° Celsius with 5% CO_2_ mixed air. The cannulated vessel was visualized on the screen with microscope connected camera and measurement system (VIA-100, Boeckeler). Endothelin-1 (ET-1, 120 pM) was used to pre-constrict the vessel for 5 minutes. Flow was applied to the vessel by changing the height of each reservoir as 55/65 cm, 50/70 cm, 40/80 cm, 30/90 cm, and 10/110 cm for 3 minutes each (Δ10, Δ20, Δ40, Δ60, and Δ100 cm H_2_O, respectively), resulting in average 60 cm H_2_O internal pressure during the experiment. After application of Δ100 cm H_2_O, papaverine (100 μM) was applied to confirm the dilatory smooth muscle function as an internal control. Flow-induced vasodilation (%) was measured by the diameter changes of vascular lumen and the equation of FIV(%) = (measured diameters – pre-constricted diameter)/(pre-pressurized inner diameter – pre-constricted diameters)*100. PA (50 μM) was applied for 1 hour prior to vessel cannulation in 30 mm dish with HEPES buffer and washed 3 times.

### Body Composition Characterization

To evaluate HFD-induced changes in body composition Kir2.1^WT^ and Kir2.1^L222I^ mice were weighed weekly beginning at 8 weeks of age, either maintained on chow or on HFD. After 14 weeks animals were subjected to nuclear magnetic resonance (NMR) for body composition analysis using the MiniSpec reader which evaluates lean, fat, and water content of the animals (56). Post-mortem fasted serum was collected for lipid panel analysis in addition to the removal of subcutaneous adipose, visceral adipose from mesentery, and brown adipose from the shoulder for weighing.

### Feeding Behavior Analysis

Feeding behavior of Kir2.1^WT^ and Kir2.1^L222I^ was evaluated using the BioDAQ Food Intake Monitor for mice (BioDAQ, Research Diets Inc., New Brunswick, NJ, USA), which allows continuous monitoring of meal patterns in undisturbed mice with minimal human interference. This is accomplished through highly sensitive scales which evaluate the weight of the food and the amount of time that the animal engages with the hopper. All the mice were habituated for 3 days to single housing and fed with chow or HFD through a BioDAQ food hopper in regular housing cages with environmental controls for one week. Mice were provided with water ad libitum (29).

### Fecal Analysis

To collect feces from Kir2.1^WT^ and Kir2.1^L222I^ mice, animals were placed individually in Technipast metabolic cages (Bugiggiate, Italy) for 48 hours with a wire bottom and a funnel system which separates urine and feces (30). Mice had access to food and water ad libitum while in the cages. Feces were collected and sent for Bomb calorimetry at the University of Michigan Metabolic Phenotyping Core to assess the caloric content of the feces in addition to the caloric content of the diet (57).

### Respiratory Exchange Rate and Indirect Calorimetry

To evaluate the respiratory exchange rate and caloric utilization (carbs compared with fat) , mice were placed in an iWorx metabolic chamber (iWorx, Inc.) in a pairwise fashion (one Kir2.1^WT^ and one Kir2.1^L222I^). The individual metabolic cage was sealed during recordings and the cage that was not being recorded was on room air. RER was measured using a GA-200 analyzer (iWorx Systems, 2013). Gas sensors were calibrated prior to experiments with gas standards containing known concentrations of O_2_, CO_2_, and N_2_ (iWorx, Inc.) (58). A gas switcher changed recording cage hourly with a one hour break at the end of each cycle, leading to RER measurements being collected every 3 hours. The ratio of VCO2/VO2 (RER) was reported using LabScribe v.4.0 (iWorx, Inc.). RER measurements were used to assess energy expenditure and to determine relative carbohydrate and lipid oxidation as a fuel source.

### Indirect calorimetry studies in Promethion system (Sable®) for metabolic profiling

Indirect calorimetry measurements were performed using the Promethion metabolic cage system (Sable Systems International, Las Vegas, NV, USA) by the Metabolic Phenotyping Core of the University of Illinois Chicago. Before data acquisition, the mice were acclimatized for 24 hours in the Promethion system and fed ad libitum with standard mouse chow and water. Food intake, energy expenditure, physical activity and substrate utilization were measured continuously 72 hours before/after diet change. Raw data were processed using ExpeData software (Sable Systems). Values are displayed as averages of the 72 hour period on each diet (chow/HFD). For energy expenditure calculation, ANCOVA analysis was applied to account for differences in bodyweight.

### Metabolomics from Mesenteric Adipose

To evaluate the metabolic pathways that may contribute to the differences in body composition between Kir2.1^WT^ and Kir2.1^L222I^ we isolated mesenteric arteries with f at that were immediately flash frozen. Tissues were then submitted to the University of Chicago Metabolomics Platform (59) for untargeted metabolomics analysis identification of individual metabolites and their relative abundance between Kir2.1^WT^ and Kir2.1^L222I^ tissues. Data are presented in heatmaps.

### Running Wheels for Activity

To evaluate the exercise activity of Kir2.1^WT^ and Kir2.1^L222I^ mice were single housed in a home cage with a running wheel attached to the top where animals had free access to run at will. This experiment was conducted over a 4 day period. Each wheel was equipped with a sensor which measured the distance run on an hourly basis and data was collected in realtime (60).

### Statistics

Results are expressed as individual data pints with the mean ± SEM or example sweeps. The normality of the data was evaluated for all data reported in this manuscript using Shapiro-Wilk tests and QQ plots. Depending on the experiment, we employed either a Student’s t-test, a one-way ANOVA, or a two-way ANOVA to determine statistical significance. When comparing two means, we employed an F-test to ascertain equal variance followed by a Student’s ttest to assess significance of the comparison. For data sets with three or more means, an appropriate type of ANOVA was employed depending on the number of independent variables. Each ANOVA analysis was coupled with Bonferroni multiple comparison testing to assess the significance between the different conditions. ANCOVA analysis was used for metabolic analysis included in Figure 6.

### Study Approval

All studies involving animals were approved by the Institutional Animal Care and Use Committee at the University of Illinois at Chicago.

## Data Availability

The supporting data values file contains all reported means and individual data points within the manuscript. Relevant information regarding the data in this manuscript will be made available from the corresponding authors upon reasonable request.

## Author Contributions

KMB, MDM, SAP, ISF, PX, and IL conceived of the study. KMB, MDM, SJA, ELM, and ISF performed experiments. KMB, MDM, SJA, and IL analyzed the data. KMB, MDM, and IL wrote the manuscript, which was edited and approved by all authors.

## Acknowledgements

Authors acknowledge all members of the Levitan and Xu labs for their insightful discussion over the work in this manuscript. We are very thankful to Dr. Sara Granados for providing initial data for the role of Kir2.1 in weight management and lipid panels. We also acknowledge Dr. Valeria Torres Irizarry for her technical assistance with metabolic studies (feeding behavior and running wheel). This research is supported by NIH T32HL144909 (KB), T32HL007829 (KB), RO1HL073965 (IL), R01HL141120 (IL), and R01HL083298 (IL).

---

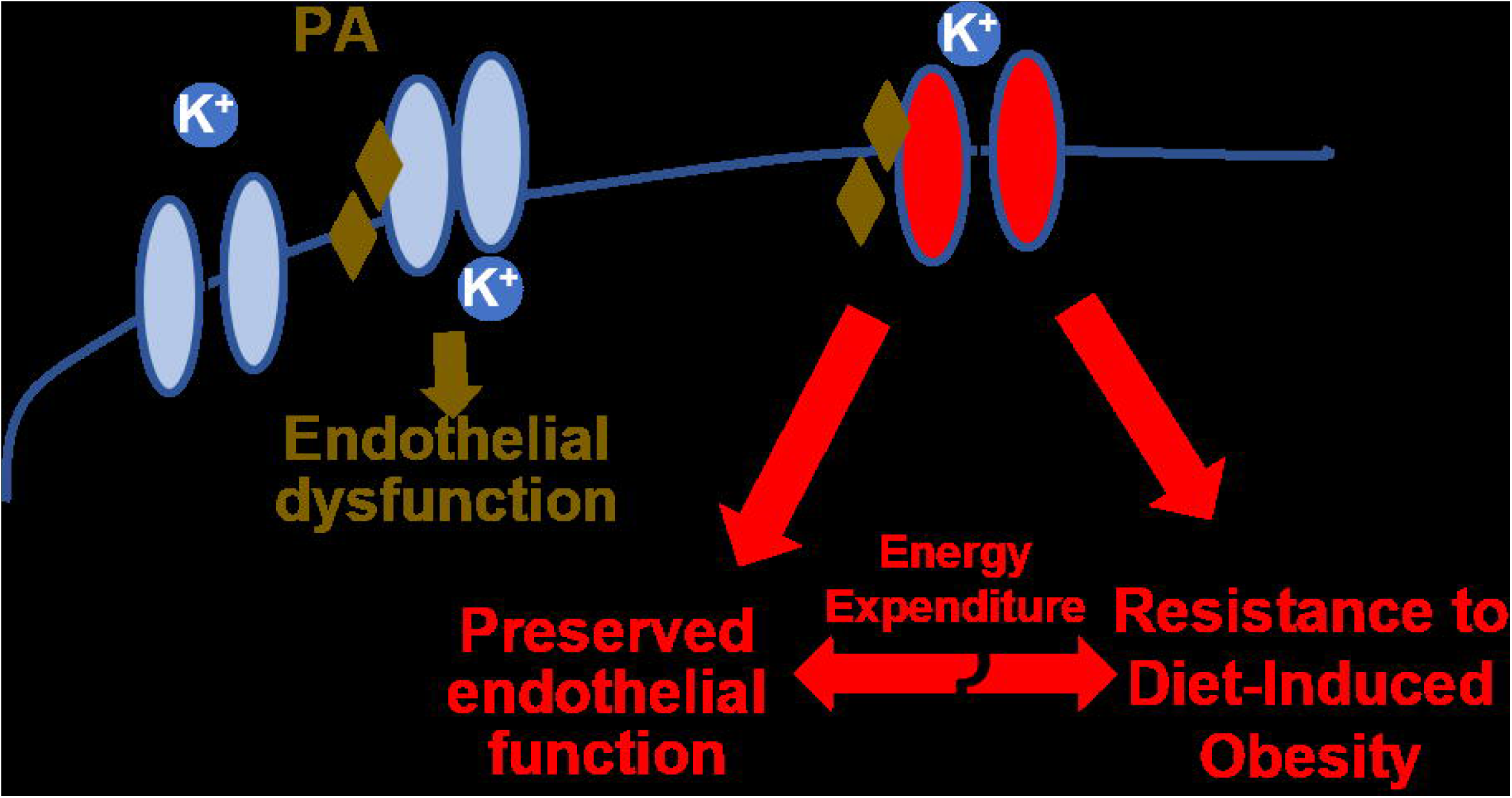

## Notes

### Competing Interest Statement

The authors have declared no competing interest.

