## Supplemental Figures for "Protection against diet-induced obesity by a single-point mutation in Kir2.1 channels"

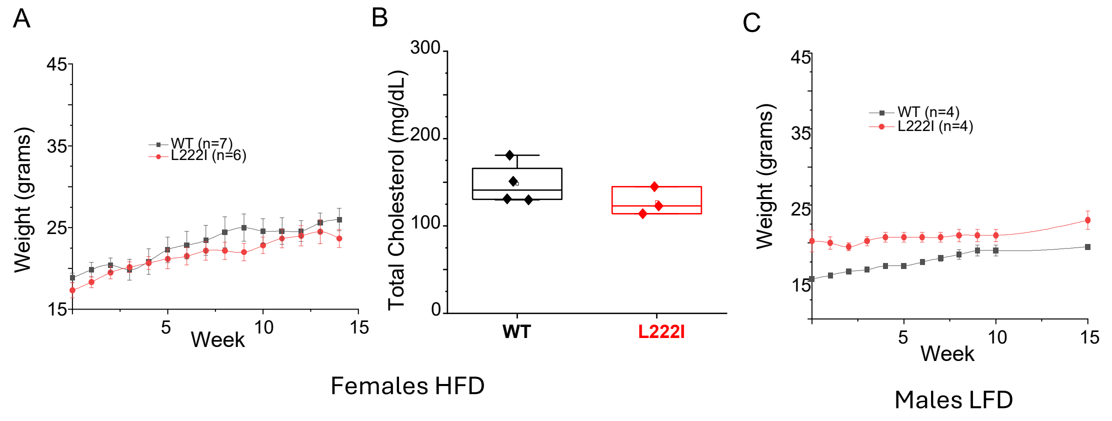


Supplemental Figure 1: Female Mice and Male Mice on LFD are unaffected by the Kir2.1$^{L222I}$ mutation. A: Time course of body weights of Kir2.1$^{WT}$ (n=7) and Kir2.1$^{L222I}$ (n=6) male mice on HFD for 14 weeks. B: Serum cholesterol levels of Kir2.1$^{WT}$ and Kir2.1$^{L222I}$ (n=3,4) female mice after 14 weeks on HFD. C: Time course of body weights of Kir2.1$^{WT}$ (n=4) and Kir2.1$^{L222I}$ (n=4) male mice on LFD for 14 weeks.

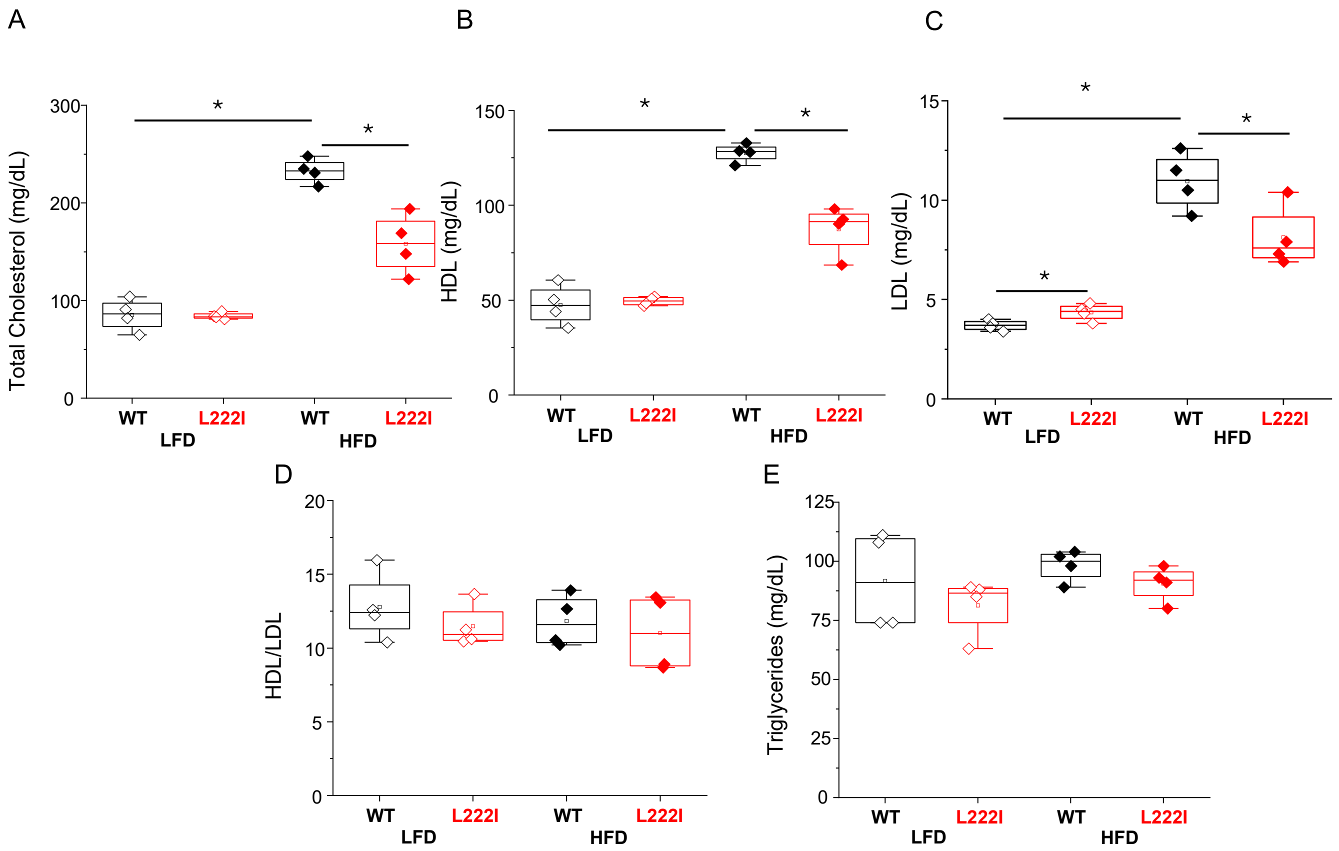


Supplemental Figure 2: Lipid Panels from Kir2.1$^{WT}$ and Kir2.1$^{L222I}$ male mice on LFD and HFD. There was no difference in lipid panels between Kir2.1$^{WT}$ and Kir2.1$^{L222I}$ on LFD except for a slight increase in LDL in Kir2.1$^{L222I}$ mice. On HFD, Kir2.1$^{WT}$ had a significant increase in cholesterol, HDL, and LDL. While there was no significant changes in Kir2.1$^{L222I}$.
